# Honeybees adjust colour preferences in response to concurrent social information from conspecifics and heterospecifics

**DOI:** 10.1101/2019.12.12.874917

**Authors:** José E Romero-González, Cwyn Solvi, Lars Chittka

## Abstract

Bees efficiently learn asocial and social cues to optimise foraging from fluctuating floral resources. However, it remains unclear how bees respond to divergent sources of social information, and whether such social cues might modify bees’ natural preferences for non-social cues (e.g. flower colour), hence affecting foraging decisions. Here, we investigated honeybees’ (Apis mellifera) inspection and choices of unfamiliar flowers based on both natural colour preferences and simultaneous foraging information from conspecifics and heterospecifics. Individual honeybees’ preferences for flowers were recorded when the reward levels of a learned flower type had declined and novel-coloured flowers were available where they would find either no social information or one conspecific and one heterospecific (Bombus terrestris), each foraging from a different coloured flower (magenta or yellow). Honeybees showed a natural preference for magenta flowers. Honeybees modified their inspection of both types of flowers in response to conspecific and heterospecific social information. The presence of either demonstrator on the less-preferred yellow flower increased honeybees’ inspection of yellow flowers. Conspecific social information influenced observers’ foraging choices of yellow flowers, thus outweighing their original preference for magenta flowers. This effect was not elicited by heterospecific social information. Our results indicate that flower colour preferences of honeybees are rapidly adjusted in response to conspecific social information, which in turn is preferred over heterospecific information, possibly favouring the transmission of adaptive foraging information within species.

---

Foraging decisions are central to an animal’s survival and reproduction; deciding where to forage in unpredictably changing environments is a major challenge that animals constantly encounter (Chittka et al. 1999; Stephens et al. 2007). Foraging decisions can be influenced in a context-dependant manner (Laland 2004; Kendal et al. 2009) by innate preferences (Lunau et al. 1996; Raine et al. 2006), previous individual experience (Sclafani 1995), and the observation or interaction with another animals at a foraging resource, i.e., social information (Heyes 1994; Leadbeater and Chittka, 2007b; Hoppitt and Laland 2013). For example, naïve individuals tend to rely more on social than individual information to gain familiarity about foraging sources (Galef and Giraldeau 2001; Galef and Laland 2005). Conversely, experienced individuals that have information on an advantageous foraging resource, will often ignore social cues. Thus, social information is only drawn upon when individual information has become outdated and acquiring up to date information may be costly (Laland 2004; Galef and Laland 2005; Kendal et al. 2009). The influence of social information on foraging decisions is taxonomically widespread (Galef and Giraldeau 2001; Valone and Templeton 2002; Grüter and Leadbeater 2014). Social information allows animals to find profitable foraging resources efficiently, instead of iteratively sampling the environment through trial and error (Galef and Laland 2005).

In most animal communities, multiple species frequently share the same foraging resources; thus members of the same (conspecific) and different (heterospecific) species can potentially act as sources and users of social information (Seppänen et al. 2007; Goodale et al. 2010; Avarguès-Weber et al. 2013; Parejo and Avilés 2016; Loukola et al. 2020). Using social information indiscriminately is not adaptive but animals should be selective when acquiring information from other individuals (Laland 2004; Kendal et al. 2018). In a multi-species context, animals have access to social information from different sources, which may give rise to a trade-off between selecting sources with a close ecological similarity, implying high competition, or sources whose ecological distance may involve less competition but a lower informative value (Seppänen et al. 2007). Furthermore, in a foraging context, social information has to be integrated with unlearned preferences (Laland and Plotkin 1993; Leadbeater and Chittka 2007a; Jones et al. 2015) and existing individual information (van Bergen et al. 2004; Jones et al. 2015) to determine foraging decisions in a context-dependant manner (Laland 2004; Kendal et al. 2009).

Social bee workers forage for nectar and pollen from rapidly changing floral resources (Heinrich 1979; Chittka et al. 1999) within multi-species communities (Fægri and van der Pijl 1979; Kevan and Baker 1983) that might offer a wide spectrum of social information. This makes bees a valuable model to explore how different sources of social information can affect learned and unlearned preferences of individuals to shape foraging decisions. Traditionally, the laboratory paradigms that investigate the use of social information in bees tend to oversimplify the real field contexts where animals naturally acquire information from others. For example, they test bees in relatively unnatural settings where a single source of social information is presented, e.g., a dead demonstrator pinned to simulated flowers (reviewed in Leadbeater and Dawson 2017). Contrastingly, bees seeking nectar and pollen in the wild might encounter far more complex circumstances where they have access to multiple sources of social information (Fægri and van der Pijl 1979; Kevan and Baker 1983) that may concur in time and space and diverge in their intrinsic relevance.

Honeybees and bumblebees of various species are in many locations sympatric and typically forage upon similar flowers due to their generalist diet (Rogers et al. 2013; Xie et al. 2016). Previous evidence indicates that bumblebees can acquire foraging information from demonstrator honeybees – in this study, the “demonstrators” were dead individuals placed on artificial flower types (Dawson and Chittka 2012). However, in more realistic conditions, a bee forager whose known floral resources have decreased in reward levels, will likely encounter other foragers, of their own and different species, feeding from different types of flowers (Fægri and van der Pijl 1979; Kevan and Baker 1983). It remains unclear how bees respond to such divergent social information, and whether in this context, social cues might modify bees’ natural preferences for particular colours in flowers (Chittka et al. 2004; Raine et al. 2006; Raine and Chittka 2007), hence influencing foraging decisions. We address these questions by testing whether honeybees might adjust their colour preferences in response to simultaneous sources of social information, i.e., a conspecific and heterospecific, each foraging from either a preferred or non-preferred flower colour.

## MATERIALS AND METHODS

### (a) Set-up

We used free-flying honeybee foragers (*Apis mellifera*) from hives located in an urban area of London, UK. Over two consecutive summers (2017-2018), we trained foragers of one hive to collect 30% sucrose solution (w/w) from a gravity feeder (von Frisch, 1965, Figure 16, p. 18), placed 2 m away from one the hive. The feeder in turn attracted foragers from the other hives, which were then included in the experiments. The feeder was refilled every day at 0800 hours, yet honeybees were also free to forage on local floral resources.

In addition, we used bumblebee foragers from two colonies (Biobest, Belgium N.V.) for the experiments. Bumblebee nests were housed in bipartite wooden nest-boxes (*l* = 29.5 × *w* = 11.5 × *h* = 9.5 cm) connected to a wooden flight arena (*l* = 77 cm, *w* = 52 cm, *h* = 30 cm) by a Plexiglas tunnel (*l* = 25 cm, 3.5 × 3.5 cm). The floor of the flight arena was covered in white laminated paper. Three plastic sliding doors located along the corridor allowed controlled access to the arena. Before and after experiments, bumblebees could freely feed upon 30% (w/w) sucrose solution from a mass feeder in the middle of the arena. Bumblebee colonies were provided with 7 g of frozen pollen (Koppert B.V., The Netherlands) every two days.

### (b) Training of bumblebees

We trained 30 bumblebee foragers (demonstrators) from two colonies (one colony per year) in a flight arena. In this group-training, foragers were allowed to enter the arena together at various times, and bumblebees learned to forage on an array of twelve plastic chips (2.4 × 2.4 cm, henceforth “flowers”) placed on the top of transparent glass vials (h = 4 cm), positioned on the floor of the arena. The array was arranged in a rectangular grid formation (3 × 4) with a separation of 7.5 cm between the edges of the flowers. We filled four yellow flowers with 10 μL of rewarding 50% (w/w) sucrose solution, and four magenta and four transparent flowers with 10 μL of unrewarding water (Figure 1A). Flowers were refilled after they were depleted by foragers and the foragers left the flower. Half of the foragers experienced the opposite colour-reinforcement contingency. Training took place for one day over 2 hours. We carried out refreshment bouts (20 min) each day prior to testing. We tracked demonstrator bees’ identity with individual number tags (Opalithplättchen, Warnholz & Bienenvoigt, Ellerau, Germany) glued to the top of the thorax by means of Loctite Super Glue Gel (Loctite, Ohio, US). The floor of the arena and flowers were cleaned with 70% ethanol after completing training.

**Figure 1.**
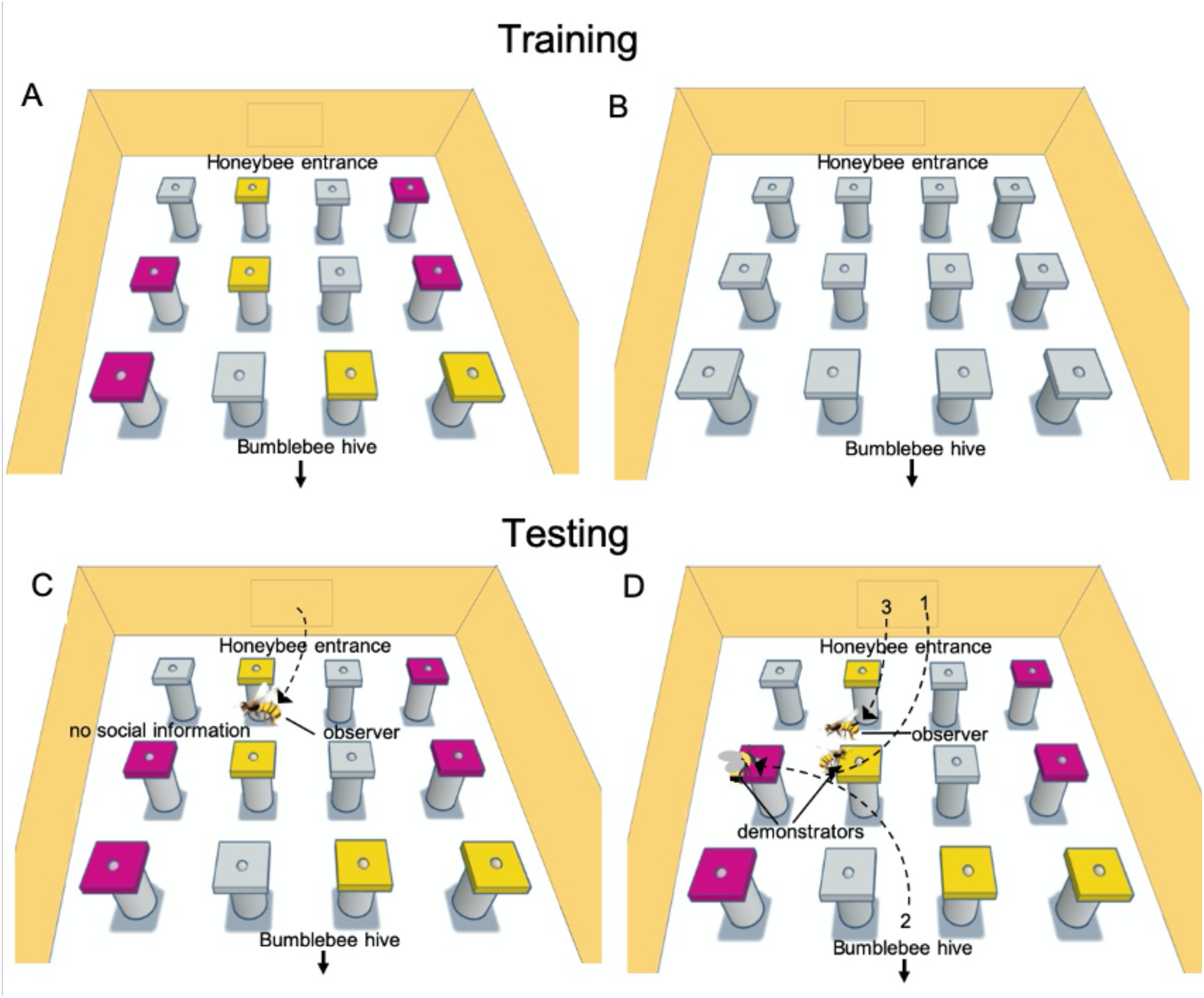
Training and testing protocols in the flight arena. (**A**) Honeybee foragers (demonstrators) were trained to find 50% sucrose solution on either four yellow or magenta flowers and water on four flowers of the alternative colour and four transparent flowers. Bumblebee foragers (demonstrators) were group trained using the same protocol (**B**) A different batch of honeybee foragers (observers) was group trained to find 50% sucrose solution on six of twelve transparent flowers and unrewarding water on the other six transparent flowers. (**C**) Honeybee observers were tested on their preparedness to forage on an unfamiliar coloured flower type, depending on its colour (yellow or magenta), once their learned flower type (transparent) yielded no reward. (**D**) Two demonstrators (one honeybee and one bumblebee) and one observer honeybee were sequentially introduced in the flight arena: (1) A honeybee demonstrator was let in the flight arena, once she foraged from a flower of her trained colour, (2) a bumblebee demonstrator was released into the arena and let to forage from a flower of her trained colour (different colour from the honeybee demonstrator), (3) an observer honeybee was then introduced to test her in a context where the flowers she previously associated with reward (transparent) were unrewarding and unfamiliar coloured flowers were demonstrated by a conspecific and a heterospecific (bumblebee) demonstrator, each foraging from non-preferred (yellow) and preferred (magenta) flowers, this design was counterbalanced.

### (c) Training of honeybees

Once we completed bumblebees’ training, we proceed to train honeybee demonstrators and observers, which took place over 18 daily sessions. Each day, we trained a set of five honeybees (demonstrators) foraging from the gravity feeder. In this group-training, honeybees learned to enter the same flight arena (located outdoors) where bumblebees were trained. We reversed the colour-reinforcement contingency so that honeybees learned to forage on flowers of the opposite colour that bumblebees were trained on. Half of the honeybees experienced four magenta flowers with 40 μL of 50% (w/w) sucrose solution, and four yellow and four transparent flowers with 40 μL of water (Figure 1A). The other half of the honeybees experienced the reverse colour-reinforcement contingency. Rewarding flowers were refilled after depletion by the foragers and the foragers left the flower. Training lasted 1 hour, consisting of six bouts (10 min). We identified trained individuals by marking their thorax with a white paint mark (Posca Pen, Worcester, UK). We cleaned the floor of the arena and flowers with 70% ethanol after completing training.

Three hours after finalising the demonstrator’s training, we selected a separate batch of five foragers (observers) to train them on a different set-up. Every day, observers were trained in group to enter the same flight arena and forage upon a rectangular grid array of twelve transparent flowers (3 × 4) (Figure 1B). To encourage foragers to sample multiple flowers during their foraging trips, only half of the flowers contained 40 μL of 50% (w/w) sucrose solution, whereas the other half contained 40 μL of water. Rewarding flowers were replenished after emptied and the forager left the flower. The positions of all the flowers were changed every bout (10 min) over 1 hour of training. We marked trained observers with a green dot to distinguish them from the demonstrators. Original honeybees trained (demonstrators) were allowed to visit the setup during this training, which did not interfere with the previous colour training, as shown in the results section. After training observers, we cleaned the flowers and the arena floor with 70% ethanol.

### (d) Testing the effect of colour preference on honeybees’ foraging decisions

We carried out this control test every day after completing training. Only two to three individuals from the batch of five observer honeybees (trained on transparent flowers) regularly showed up at the setup at the time of testing. Thus, we only tested two daily individuals. Overall, we tested 20 honeybees individually (control group) in a context where transparent flowers were unrewarding. We assessed whether they might show a natural preference to inspect and forage from one flower type between two unfamiliar alternatives, i.e., yellow and magenta. One honeybee was let in the arena to explore a rectangular grid array of twelve flowers (section b, above), consisting of four familiar transparent flowers containing 20 μL of water and eight unfamiliar flowers, four yellow and four magenta, all filled with a scentless reward of 20 μL of 50% (w/w) sucrose solution, (Figure 1D). The test began once the individual inspected any flower. That is, the honeybee approached the flower by displaying a slow side-to-side hover, with its head oriented towards it and within at least one body length (Ings et al. 2009). The test concluded once the honeybee landed and foraged upon any unfamiliar flower, or 3 minutes after the test started. As this test was designed to evaluate the influence of honeybees’ colour preferences on an actual foraging decision, rather than measuring their innate colour preferences, we regarded the flower type where the individual landed and foraged as preferred over the alternative type. To prevent re-testing the same individuals, we captured honeybees, after concluding the test, to give them a distinctive red paint mark. The flowers and arena floor were cleaned with 70% ethanol between tests. To evaluate honeybees’ inspection of flowers before they chose a flower to forage (foraging decision), we recorded the test with a sport camera (Yi, Xiaomi Inc. China) featuring a recording frame rate of 30 fps and a resolution of 720 p (1,280 × 720 pixels). The camera was positioned 20 cm above the entrance of the bumblebee nest (Figure 1). Its field of view was adjusted such that it looked down into the arena at ∼50° from a horizontal angle.

### (e) Testing the influence of social information on foraging decisions

To evaluate the influence of simultaneous sources (conspecific and heterospecific) of social information on honeybee’s inspection of either yellow or magenta flowers and their subsequent foraging decisions, we tested 45 honeybees in the same context as individuals in the control group (section d) but in the presence of one bumblebee and one honeybee demonstrator, each foraging upon either a yellow or magenta flower. We introduced the demonstrators and observer in the arena in the following order: 1) a honeybee demonstrator was let in through a sliding door on the back wall of the arena (Figure 1D). Once she began to feed from a flower of her trained colour, 2) a bumblebee demonstrator was released from the Plexiglas tunnel that connected the arena to the nest-box (Figure 1D). When she started to forage from a flower of her trained colour (opposite colour as the honeybee demonstrator), 3) a honeybee observer was let into the arena through the sliding door on the back wall. Due to training, both demonstrators swiftly landed and foraged exclusively on flowers of their trained colour. We tested 19 observer honeybees with a conspecific demonstrator foraging on magenta flowers and a heterospectific foraging on yellow flowers and 26 observers with the reversed colour-reinforcement contingency.

The test began once the observer honeybee inspected (see section d) any occupied or unoccupied flower. The test concluded once the observer landed and foraged upon any unfamiliar flower, or 3 minutes after the test started. In the test, demonstrators moved freely between flowers after depleting them, as they naturally do within inflorescences. Thus, we only considered that observers foraged on an unfamiliar flower when they actually fed upon the sucrose solution reward from unemptied flowers. To prevent re-testing the same individuals, we caught tested honeybees and marked them with a distinctive red paint mark. The flowers and arena floor were cleaned with 70% ethanol between tests. To evaluate honeybees’ inspection of flowers and their interactions with the demonstrators before they made a foraging decision, the test was recorded as described in section (d) above.

### (d) Analyses

We analysed the behaviour of tested honeybees from video recordings, using the BORIS behavioural observation software (Friard and Gamba, 2016). To assess individuals’ inspection of either yellow or magenta flowers and their preference to forage upon one type of flower over the other, we analysed two main behavioural categories. That is, the frequency of inspecting transparent, yellow and magenta flowers (Ings et al. 2009; Balamurali et al. 2018) and individual’s foraging choice, i.e., the yellow or magenta flower where individuals landed and foraged. We used logistic analysis to explore the influence of flower colour on the likelihood of honeybees inspecting a flower for the first time, the likelihood of foraging on a flower type, as well as the proportion of transparent, yellow and magenta flowers they inspected. For the latter two variables, we took underdispersion into account via a quasinomial model.

For honeybees exposed to social information, we considered inspection of occupied flowers as an indicator that observers detected the demonstrator’s presence. Thus, four different scenarios were possible before honeybees made a foraging decision: they could have detected both demonstrators, either the honeybee or bumblebee demonstrator, or none of them. We compared the likelihood of each situation against the expected probability with a Chi-Square Goodness of Fit Test. Nine individuals that did not detect the presence of the demonstrators were not considered for all analyses.

We used logistic analysis to determine whether colour preferences and concurrent conspecific and heterospecific social information had a similar influence on foraging decisions of honeybees. Social information was included as a predictor variable (binary: 1= present, 0= absent) for the likelihood of honeybees landing on either an unfamiliar yellow or magenta flower.

Further, to evaluate the influence of social information on honeybees’ readiness to forage on an unfamiliar flower, we compared two measurements between the control group and the group exposed to social information. That is, the total number of flowers that individuals inspected before they chose a flower to forage, and the time it took them to make such decision (latency to forage). We analysed both measurements to explore whether observers’ readiness to forage was influenced by the first foraging demonstrator they detected in the test (i.e., honeybee or bumblebee). This was irrespective of whether they detected only one or both demonstrators. These analyses were conducted with a Wilcoxon rank sum test.

To determine whether observers that detected both demonstrators inspected the flowers occupied by a honeybee and bumblebee at a similar frequency, we compared, with a Wilcoxon signed rank test, the proportion of flowers occupied by each demonstrator that observers inspected. We also analysed, with a Wilcoxon rank sum test, whether honeybees that only detected either a conspecific or heterospecific demonstrator, differed in the proportion of occupied flowers that they inspected, relative to all the flowers they inspected during the test. We used logistic analysis to explore the influence of flower colour and demonstrator’s species on the likelihood of honeybee observers selecting the demonstrated type of flower and the likelihood that observers would forage on a flower occupied by either a honeybee or bumblebee, after approaching it for the first time. For the latter variable, we took underdispersion into account via a quasinomial model.

The instances in which observers inspected an occupied flower that did not progress into landing and foraging were recorded as rejections. We evaluated whether observers rejected the flowers occupied by a honeybee or bumblebee demonstrator at a similar frequency. For observers that detected both demonstrators, we compared the proportion of times that they rejected the flowers occupied by each demonstrator, using a Wilcoxon signed rank test. We also compared, with a Wilcoxon rank sum test, the total number of occupied flowers that observers rejected when they only detected either the honeybee or bumblebee demonstrator.

To assess whether observers altered their inspection of flowers after they initially detected the presence of either a foraging honeybee or bumblebee, we adjusted the frequency of observers’ inspection of flowers by normalising the data through an index calculation, with the following equation:

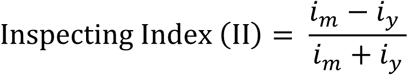

where, *i*_*m*_ and *i*_*y*_ are the frequency that the observer inspected magenta and yellow flowers, respectively.

In the indices, a negative value (minimum -1) equates to a preference to inspect yellow flowers, whereas a positive value (maximum +1) equates to a preference to inspect magenta flowers, an index near equal to zero implies that the observers either inspected both types of flowers equally, or they did not inspect the flowers at all. We calculated inspecting indices (II) based on the sequence of events that preceded the honeybees’ foraging decisions. That is, each inspecting index (II) represented observers’ inspection of flowers, before and after they detected either demonstrator (honeybee or bumblebee) foraging on a flower. We compared indices against chance expectation (Index = 0) with a Wilcoxon signed-rank test. A significant switch from a negative or positive value, before the observer detected a demonstrator, to a positive or negative value, after this occurred, indicates that observers modified their inspection of yellow or magenta flowers in response to conspecific or heterospecific social information. All analyses were conducted using R statistical software (R Core Team, 2019).

### Ethical Note

Bees were kept in their natural colony environment in unaltered dark conditions. The gravity feeder where honeybees foraged from, was only filled when no individuals were present to avoid disturbance. Bumblebees were fed with a minimum disturbance under red light, which is poorly visible to bees. Colonies were not food deprived during experiments. Only bee foragers that freely engaged in foraging behaviour were trained and tested. Honeybees were tagged with a dot of paint on the thorax while feeding and bumblebees were carefully handled with dissection forceps to glue number tags on their thoraxes.

## RESULTS

### Effect of colour preference on honeybees’ foraging decisions

In a context where a familiar flower type ceases to yield reward, how does colour preference influence honeybees’ exploration of contiguous, novel flowers and ultimate move to foraging upon a new flower type? We carried out a control test to assess whether honeybees preferentially inspect one unfamiliar flower type over another, and whether this might affect their ultimate foraging decision (move to foraging upon a new flower type). We trained honeybees (observers) to find either a food reward (50% sucrose solution) or unrewarding water on transparent flowers (Materials and Methods). We then we tested bees in a context where transparent flowers were unrewarding and two unfamiliar yellow and magenta flower types delivered a food reward (scentless 50% sucrose solution). The majority of individuals (77%) did not inspect the familiar transparent flowers at all; rather honeybees preferred to inspect and forage on magenta flowers. The proportion of flowers inspected by honeybees was influenced by flower colour (Logistic regression: *F* = 35.86, *N* = 20, P < 0.001; Fig. 3.2A). Flower colour also influenced the likelihood of honeybees first exploring a flower type (Logistic regression: *χ*^*2*^_1_ = 15.44, *N* = 20, P < 0.001; Fig.3.2B) and the likelihood of selecting a flower to forage (Logistic regression: *F* = 30.04, *N* = 20, P < 0.001; Fig. 3.2C). This shows that honeybees had a colour preference for magenta over yellow flowers.

**Figure 3.2.**
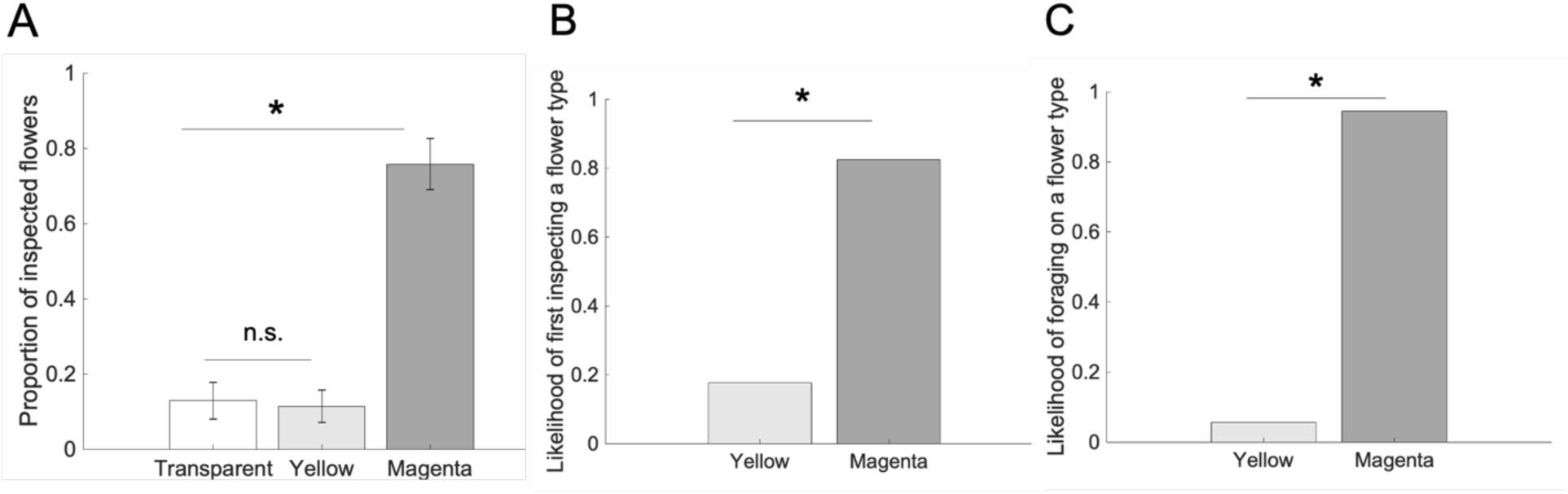
Honeybees prefer magenta over yellow flowers **(A)** Honeybees inspected magenta flowers at a higher proportion than yellow and transparent flowers. **(B)** The likelihood of honeybees inspecting an unfamiliar flower for the first time was higher for magenta compared to yellow flowers. **(C)** A higher proportion of honeybees preferred to forage on magenta than yellow flowers. Bars = mean. Horizontal lines = standard error. Asterisks = p < 0.05.

### Influence of social information on honeybees’ foraging decisions

In a field-like context, including simultaneous and divergent conspecific and heterospecific social information, we investigated how this social information might affect honeybees’ inspection of flower types and ultimate foraging decisions. We considered that observers detected the demonstrators’ presence once they inspected an occupied flower for the first time. We found no difference in the proportion of observers that detected both demonstrators, only the honeybee or bumblebee demonstrator, or none of them (Chi-Square Goodness of Fit Test: *χ*^*2*^_3_ = 2.2, *N* = 45, *P* = 0.31; Fig. 3A). The likelihood of foraging on an unfamiliar flower type was not influenced by the presence of social information (Logistic regression: *χ*^*2*^_1_ = 0.9, *N* = 56, *P* = 0.34; Fig. 3.2B). Similar to the control group, the vast majority of observers (90%) did not inspect the transparent flowers at all. These results suggest that both individual colour preference and social information similarly influenced honeybees when locating a new foraging resource. Compared to observer honeybees that first detected a foraging bumblebee in the test, individuals in the control group, whose foraging decisions were solely influenced by colour preference, made faster choices (Wilcoxon rank-sum: *W* = 71, *N* = 35, P = 0.007; Fig. 3.3C) and inspected fewer flowers (Wilcoxon rank-sum: *W* = 77, *N* = 35, P = 0.011; Fig. 3.3D). However there was no difference between the control group and those observers that first detected the presence of a foraging conspecific (Wilcoxon rank-sum: latency: *W* = 133, *N* = 30, *P* = 0.29; Fig. 3.3C; number of flowers: *W* = 125, *N* = 30, *P* = 0.45; Fig. 3.3D). Further, when observers first detected the conspecific demonstrator, they made faster foraging decisions (Wilcoxon rank-sum: *W* = 184, *N* =29, P < 0.001; Fig. 3.3C), preceded by less inspection of flowers (Wilcoxon rank-sum: *W* = 176.5, *N* = 29, P < 0.001; Fig. 3.3D), than when they first detected the heterospecific bumblebee. These results suggest that both individual preference for magenta flowers and conspecific social information similarly affected honeybees’ foraging choices, enabling swift decisions. Conversely, observers that initially detected a foraging heterospecific, explored the flowers extensively before making a foraging decision.

**Figure 3.**
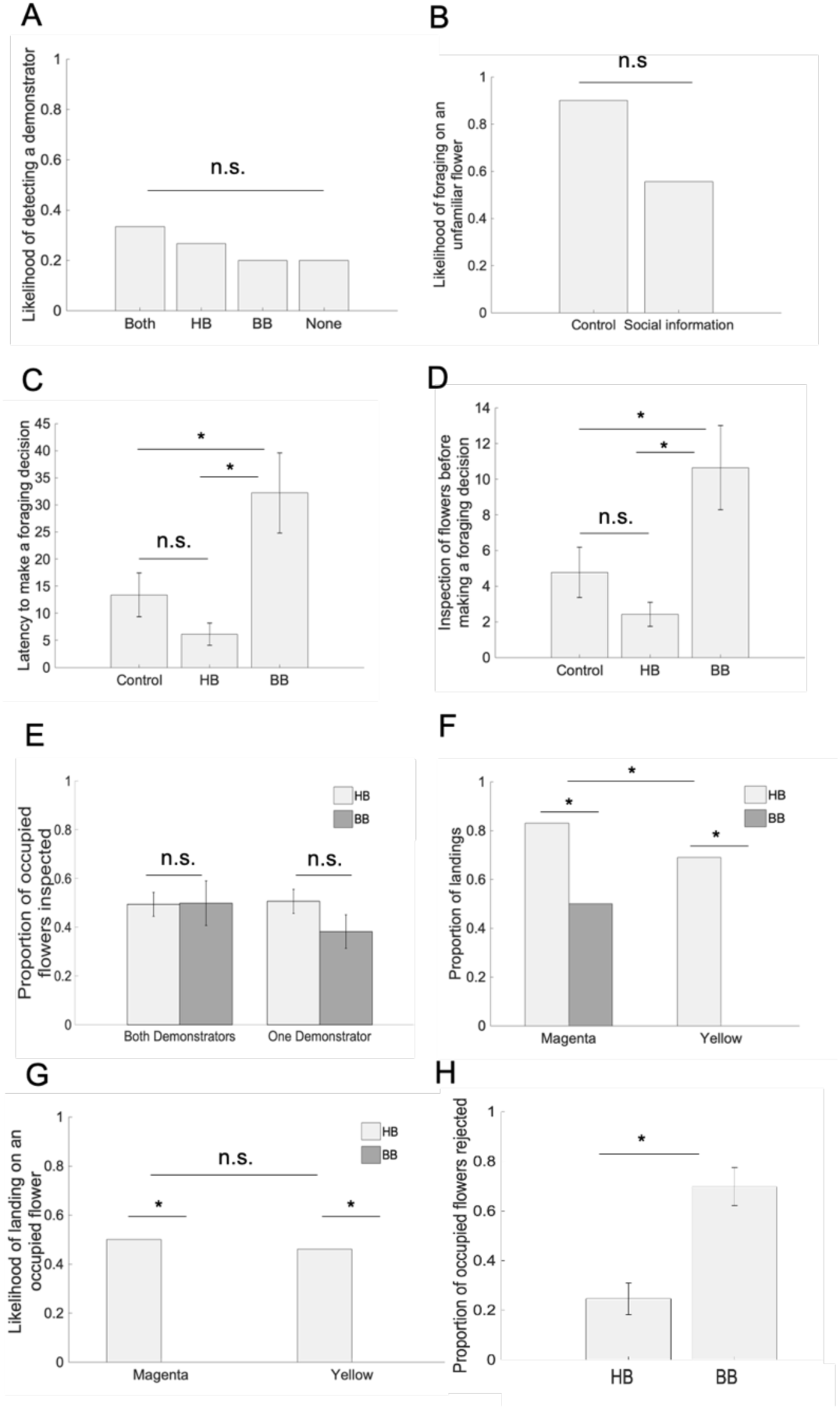
Concurrent conspecific and heterospecific social information influences foraging behaviour of honeybees **(A)** Observer honeybees were equally likely to detect both demonstrators, only the honeybee or bumblebee demonstrator, and none of them. **(B)** The proportion of honeybees that foraged on unfamiliar flowers was not different between observers that detected the presence of demonstrators and individuals in the control group (no social information). **(C)** Honeybees in the control group and those that first detected a foraging conspecific made faster foraging decisions than observers that first detected a foraging heterospecific. **(D)** Honeybees in the control group and those that first detected a foraging conspecific inspected fewer flowers before making a foraging decision than observers that first detected the presence of a foraging heterospecific. **(E)** Honeybee observers that detected only one or both demonstrators during the test; similarly inspected flowers occupied by the honeybee and bumblebee demonstrator. **(F)** Demonstrators’ species and flower colour had an effect on the proportion of honeybees that foraged on unfamiliar flowers, yet no interaction of main effects was found. **(G)** Once observers detected a conspecific demonstrator foraging on an unfamiliar flower type, they were more likely to land and forage on such a flower, but they never foraged on a flower occupied by a heterospecific demonstrator. **(H)** Observers rejected flowers occupied by a bumblebee at a higher proportion than flowers occupied by a conspecific. HB = honeybee, BB = bumblebee. Bars = mean. Horizontal lines = standard error. Asterisks = p < 0.05. n.s. = not significantly different.

Honeybee observers that detected either one or both demonstrators, similarly inspected flowers occupied by either the honeybee or bumblebee demonstrator (Wilcoxon rank-sum; one demonstrator: *W* = 34.5, *N* =16; *P* = 0.45; both demonstrators: Wilcoxon signed-rank: *V* = 22.5, *N* = 13, *P* = 1; Fig. 3.3E). However, the likelihood that an observer would forage on an unfamiliar flower type was influenced by the demonstrators’ species (Logistic regression: *χ*^2^_1_ = 11.78, *N* = 45, P < 0.001; Fig. 3.3F) and flower colour (Logistic regression: *χ*^2^_1_ = 5.08, *N* = 45, P = 0.024; Fig. 3.3F) but was only marginally influenced by the statistical interaction between these main effects (Logistic regression: *χ*^2^_1_ = 3.17, *N* = 45, *P* = 0.07; Fig. 3.3F). Further, demonstrators’ species influenced the likelihood of observers foraging on an occupied flower after approaching it for the first time (Logistic regression: *χ*^2^_1_ = 20.51, *N* = 51, P < 0.001; Fig. 3.3G). That is, observers readily responded to the presence of a foraging conspecific by joining such demonstrator on the unfamiliar flower type. This response was not affected by flower colour (Logistic regression: *χ*^2^_1_ = 0.03, *N* = 51, *P* = 0.82; Fig. 3.3G) nor by any statistical interaction between main effects (Logistic regression: *χ*^2^_1_ = 0, *N* = 51, *P* = 1; Fig. 3.3G). In contrast, observer honeybees rejected the flowers occupied by a bumblebee demonstrator at a higher proportion (Wilcoxon signed-rank: *V* = 3, *N* = 13, P = 0.008; Fig. 3.3H) than the flowers occupied by a conspecific.

### Adjustment of honeybees’ colour preference in response to social information

The inspecting indexes (II), described in (d), allowed us to analyse honeybees’ inspection of flowers as a flexible process that culminated in a foraging decision, and whose variation in response to social information was measurable in terms of an index value. Observer honeybees showed no preference to inspect magenta or yellow flowers before they detected the bumblebee demonstrator foraging on either a magenta (Wilcoxon signed-rank test: *V* = 20, *N =* 9, *P* = 0.15; Fig.4A) or yellow flower (Wilcoxon signed-rank test: *V* = 12, *N* = 9, *P* = 0.11; Fig. 4A). After honeybees detected a bumblebee foraging from a magenta flower, they preferentially inspected this type of flower (Wilcoxon signed-rank: *V* = 26, *N* = 9, P = 0.026; Fig. 3.4A). This preference did not occur after they observed a bumblebee demonstrator foraging on a yellow flower, and instead they showed no preference to inspect either type of flower (Wilcoxon signed-rank: *V* = 18.5, *N* = 9, *P* = 0.5; Fig. 3.4A). Honeybees showed no preference to inspect either type of flower before they detected a conspecific foraging upon either a magenta (Wilcoxon signed-rank: *V* = 16.5, *N* = 11, *P* = 0.61; Fig. 3.4B) or yellow flower (Wilcoxon signed-rank: *V* = 23, *N* = 11, *P* = 0.26; Fig. 3.4B). However, observer honeybees did respond to the presence of a conspecific foraging on either flower type by increasing their inspection of demonstrated flowers (Wilcoxon signed-rank: magenta: *V* = 52, *N* = 11, P = 0.006; yellow: *V* = 1, *N* = 13, P = 0.002; Fig. 4B). These results show that honeybees’ inspection of unfamiliar flowers underwent a sequential adjustment in response to conspecific and heterospecific social information, which ultimately affected their foraging decisions.

**Figure 4.**
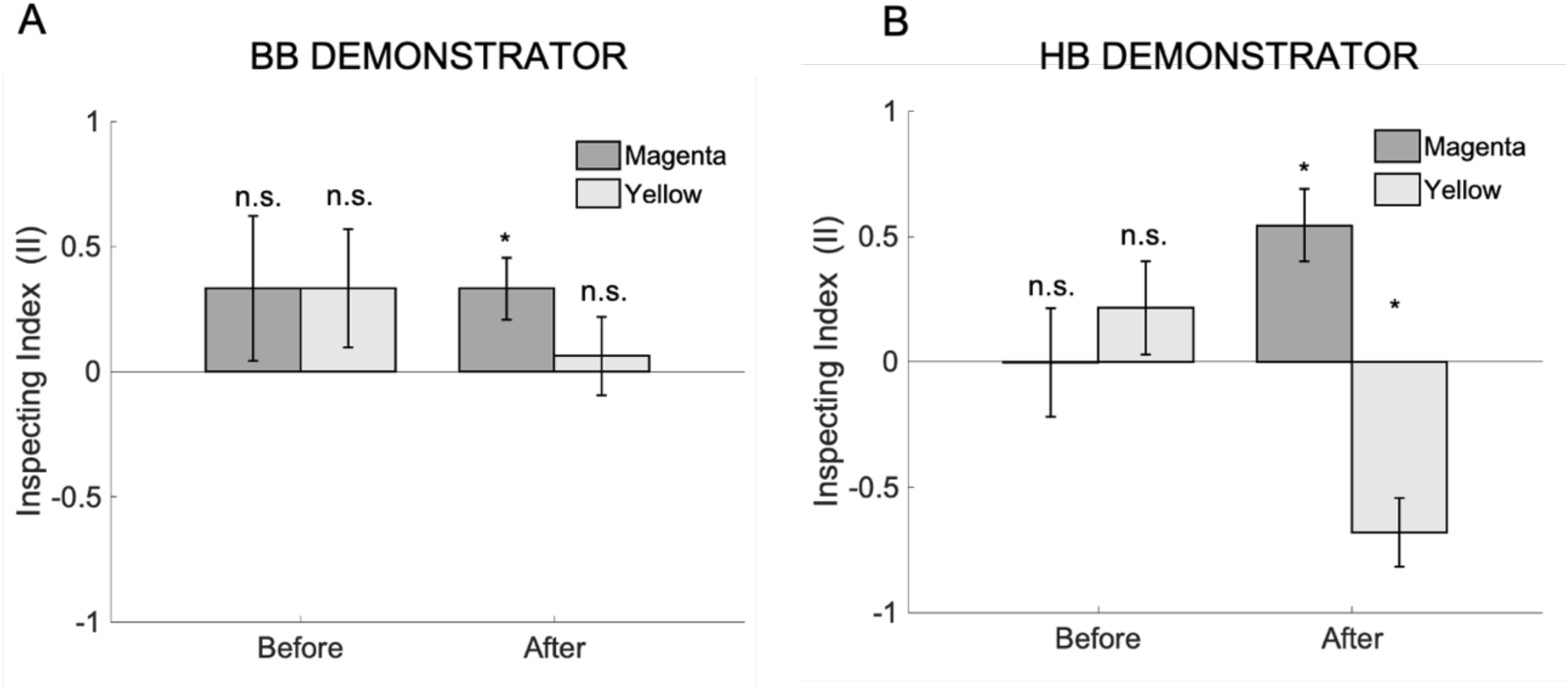
Honeybees adjust their exploration of flowers and foraging preferences in response to concurrent conspecific and heterospecific social information. Inspecting indices (II) compare to chance level (Index = 0) the inspection of magenta (dark grey bars) and yellow (light grey bars) flowers that honeybee observers had before and after they detected either a conspecific or heterospecific demonstrator foraging on either type of flower. **(A)** Before observers detected a bumblebee demonstrator foraging on either a magenta or yellow flower, they showed no preference to inspect neither of these flower types. After observers detected a bumblebee demonstrator foraging on a magenta flower, they inspected this type of flowers more frequently. However, when observers detected the bumblebee demonstrator foraging on a yellow flower, this did not affect their inspection of yellow or magenta flowers. **(B)** Before observers detected a conspecific foraging upon either a yellow or magenta flower, they showed no preference to inspect magenta or yellow flowers. After observers detected a conspecific demonstrator foraging on either a yellow or magenta flower, they increased their inspection of the demonstrated type of flower. HB = honeybee, BB = bumblebee. Index = 0. Bars = mean. Horizontal lines = standard error. Asterisks = p < 0.05. n.s. = not significantly different between groups or from chance level.

## DISCUSSION

The results presented here provide evidence that honeybees’ colour preferences can be adjusted in response to simultaneous social information from conspecifics and heterospecifics. Thus, specific social information differently influences honeybees’ exploration of flowers and consequent foraging decisions. Unexpectedly, honeybees payed little attention to the transparent flowers on which they were trained. Instead, prior to making a foraging decision, they inspected more frequently the magenta flowers (control group) or the type of flower demonstrated by a conspecific.

Even though inspecting flowers from a close distance does not provide foragers with information on the reward status of a flower, it is important in the process of choosing a flower (Lunau et al. 1996). In the absence of social information, honeybees’ natural preference for magenta flowers influenced inspection and choices of flowers. Magenta is actually a mixture of blue and red, since the red component is not fully perceived by bees (Menzel and Shmida 1993), they perceive magenta flowers as blue (Chittka and Waser, 1997; Waser and Chittka 1998). Honeybees and bumblebees usually have innate biases for colours in the violet to blue range of the spectrum (Giurfa et al. 1995; Chittka et al. 2004), which adaptively correlates with the nectar production of local flowers (Giurfa et al. 1995; Chittka et al. 2004). Although colour biases may be overridden by individual learning (Raine et al., 2006), they potentially govern foraging decisions in bees when selecting among novel flower types (Gumbert 2000). In the absence of social information, flower choices of tested honeybees conceivably resulted from either their innate colour preferences or previous experience in the field with local flower species as both factors can be linked with the likelihood of finding a profitable food reward (Giurfa et al. 1995; Chittka et al. 2004).

In the wild, honeybee foragers are likely to encounter members of the same and different species foraging concurrently in a flower patch including distinct flower types (Fægri and van der Pijl 1979; Kevan and Baker 1983). Our results demonstrate that such social information is integrated with honeybees’ colour preferences during the process of making a foraging decision. In this realistic scenario, with two simultaneous sources of conspecific and heterospecific social information, when the first foraging demonstrator that honeybees detected was a member of the same species, they promptly responded to social information by joining the demonstrator on the unfamiliar flower type. This evidence is consistent with previous findings in bumblebees (reviewed in Leadbeater and Dawson 2017) and supports the notion that the use of conspecific social information in bees may be mediated by local enhancement, with foragers being attracted to flowers occupied by members of the same species (Leadbeater and Chittka 2007a). Such a response potentially optimises the foraging efficiency of bees by facilitating their movements between different flower types (Leadbeater and Chittka 2005; Kawaguchi et al. 2007).

Honeybees and bumblebees often forage upon similar floral resources; it is then conceivable that they may concur in the same flower patches (Rogers et al. 2013; Xie et al. 2016). In our experiments, we used freely moving bumblebee demonstrators to elucidate how social information might flow between these species. Our results indicate that in the likely scenario of a honeybee exploring a flower patch and encountering both a conspecific and heterospecific (bumblebee) individual foraging from different flower types, its readiness to use social information and make a foraging decision might depend on the first observed species providing social information. That is, observer honeybees that detected a conspecific demonstrator before a heterospecific bumblebee made swifter foraging decisions than those observers that first detected the bumblebee demonstrator, which spent more time exploring the flowers before making a foraging decision. This evidence suggests that in a foraging context, honeybees’ discrimination of conspecifics and heterospecifics might determine the selective use of social information. Theory predicts that using social information from heterospecifics should be favourable when there is a large niche overlap (Seppänen et al. 2007) yet, on adaptive grounds, when animals can select between social information from conspecifics and heterospecifics, they should typically favour the former as this naturally reflects its ecological needs (Jaakkonen et al. 2015). In line with this, honeybees consistently selected the type of flowers demonstrated by conspecifics, even though they attended to the presence of both demonstrators whilst exploring the set-up. Prioritising conspecific over heterospecific choices can be explained due to its high fitness value (Seppänen et al. 2007; Jaakkonen et al. 2015) but it can also serve to the transmission of novel behavioural traits or preferences that may be adaptively valuable for a particular species (Laland and Plotkin 1993; Jaakkonen et al. 2015; Alem et al. 2016; Danchin et al. 2018).

Floral reward levels differ strongly among plant species and constantly change over time in an unpredictable manner (Heinrich 1979). To achieve efficient foraging, bees can rapidly learn to associate floral traits such as colour, shape and scent with reward quality in flowers (Chittka et al. 1999). Bees can be initially attracted to forage from an unfamiliar flower species via either innate and learned colour preferences (Giurfa et al. 1995; Gumbert 2000; Chittka et al. 2004; Raine and Chittka 2007) or using social information (Leadbeater and Chittka 2007b; Grüter and Leadbeater 2014; Leadbeater and Dawson 2017). In the wild, bees seeking floral resources are unlikely to experience asocial and social cues separately, rather they may frequently be exposed to a complex combination of cues potentially affecting their flower choices. Our findings demonstrate that simultaneous conspecific and heterospecific social information influences honeybees’ colour preferences, which may in turn shape the process of acquiring new information underlying foraging decisions.

Compared to honeybees in the control group (i.e., no social information), the exploration behaviour of honeybees that observed a conspecific or heterospecific demonstrator foraging upon the less preferred yellow flowers, reflected a more evenly distributed inspection of magenta and yellow flowers. That is, the presence of either demonstrator on a yellow flower increased the ‘attractiveness’ of such flower type for honeybees, possibly via stimulus enhancement, an effect widely described in social learning literature (Heyes 2012). Remarkably, honeybees’ inspection of flowers, naturally biased towards magenta flowers, underwent a sequential and flexible adjustment prior to make a foraging decision. Such adjustment was modulated by visual foraging information from members of the same and different species.

It has been demonstrated that bees are attracted to the presence of foraging conspecifics when presented with unfamiliar flowers which may lead them to identify new rewarding flower species (Leadbeater and Chittka 2007a; Jones et al. 2015). Our results indicate that the presence of a foraging conspecific not only influenced honeybee observers to select flowers that matched their colour preference (magenta) but such conspecific social information also outweighed observers’ colour preference so that they selected the normally non-preferred yellow flowers. Whereas intraspecific social transmission of foraging information may be more stable when it reinforces a prior preference (Laland and Plotkin 1990), such preferences may be adjusted and potentially overridden in response to conspecific social information about a novel foraging resource (Jones et al. 2015). This may enable bees to reasonably adapt to wildly varying floral reward levels. Thus, if the presence of a foraging conspecific can reliably be associated with a rewarding outcome, the use of such social cue should be reinforced to consistently influence bees’ flowers choices across foraging contexts (Leadbeater and Chittka 2009). Interestingly, in our experiments the presence of a foraging heterospecific also increased honeybees’ attraction to the demonstrated flower types, yet heterospecific social information did not influence honeybees’ foraging choices. In fact, honeybee observers never landed on the flowers occupied by the bumblebee demonstrator, in contrast to the flowers occupied by the conspecific demonstrator. Even though, heterospecific social information is predictably valuable when there is a large niche overlap (Seppänen et al. 2007), conspecifics choices might offer a more predictable social cue, decreasing the risk of acquiring maladaptive information (Giraldeau et al. 2002) as conspecifics share the same ecological needs (Seppänen et al. 2007; Goodale et al. 2010).

Multiple social and asocial cues can potentially affect animals’ foraging decisions in different circumstances (Sclafani 1995; Galef and Giraldeau 2001; Grüter and Leadbeater 2014). The effect of social information on previous, innate or learned preferences influences the foraging decision of a particular individual and may also promote the transmission of adaptive information about food sources (Laland and Plotkin 1990; Galef and Giraldeau 2001). Despite the fact that social information offers a clear advantage to individuals exploring novel foraging resources (Galef and Giraldeau 2001), its use should not be indiscriminate but respond to particular circumstances in order to lead to adaptive choices (Laland 2004; Kendal et al. 2018). Our results extend previous evidence showing that conspecific social information is commonly used in situations of uncertainty (Kendal et al. 2004; van Bergen Yfke et al. 2004; Galef et al. 2008; Smolla et al. 2016), such that it can outweigh both predetermined individual preferences (Dugatkin 1996; Jones et al. 2015) and heterospecific social information (Jaakkonen et al. 2015) to influence ecologically relevant decisions in animals.

It is conceivable that natural selection should favour the salience of social stimuli with high ecological relevance (Seppänen et al. 2007; Leadbeater and Dawson 2017), conspecifics in our experiments. Whether such salience can similarly operate to select information from heterospecific sources, based on their relative informative value, deserves further consideration. We provide an ecologically relevant picture of the process thereby multiple non-social and social cues may shape foraging decisions of bees. Thus, findings presented here contribute to our understanding of the flexibility of individual preferences and their adjustment in response to different sources of social information. Our results in turn shed light on the selectivity of animals for conspecific over heterospecific information as a possible mechanism to facilitate the social transmission of foraging information within species.

